# Resting-State Functional Networks of Different Topographic Representations in the Somatosensory Cortex of Macaque Monkeys and Humans

**DOI:** 10.1101/775569

**Authors:** John Thomas, Dixit Sharma, Sounak Mohanta, Neeraj Jain

## Abstract

Information processing in the brain is mediated through a complex functional network architecture whose comprising nodes integrate and segregate themselves on different timescales. To gain an understanding of the network function it is imperative to identify and understand the network structure with respect to the underlying anatomical connectivity and the topographic organization. Here we show that the previously described resting-state network for the somatosensory area 3b comprises of distinct networks that are characteristic for different topographic representations. Seed-based resting-state functional connectivity analysis in macaque monkeys and humans using BOLD-fMRI signals from the face, the hand and rest of the medial somatosensory representations of area 3b revealed different correlation patterns. Both monkeys and humans have many similarities in the connectivity networks, although the networks are more complex in humans with many more nodes. In both the species face area network has the highest ipsilateral and contralateral connectivity, which included areas 3b and 4, and ventral premotor area. The area 3b hand network included ipsilateral hand representation in area 4.

The emergent functional network structures largely reflect the known anatomical connectivity. Our results show that different body part representations in area 3b have independent functional networks perhaps reflecting differences in the behavioral use of different body parts. The results also show that large cortical areas if considered together, do not give a complete and accurate picture of the network architecture.

**Highlights:**

- Somatosensory resting-state functional network is not uniform across the entire area 3b. Different body part representations have different connectivity networks.
- These functional connectivity networks have many similarities in the two primate species, i.e. macaque monkeys and humans, although the human network is more complex.
- In both the species network of the face representation is most extensive, which includes ipsilateral face motor cortex and PMv in both hemispheres.
- The hand representation in area 3b has connectivity with ipsilateral hand motor cortex.
- Bilateral connectivity with homologous and nonhomologous area 3b representations was observed only in humans.
- The functional connectivity networks largely reflect the underlying anatomical connectivity.

## 1. INTRODUCTION

Since early descriptions of the cortical areas based on cytoarchitecture (Brodmann, 1909), various anatomical, neurophysiological and neuroimaging studies have given new insights into functional segregation across brain regions (see Amunts and Zilles, 2015). Cortical parcellation combining multiple features viz. cyto-, myelo- and immuno-architecture, topography, function and connectivity analysis help better define areal and sub-areal boundaries in the cortex and understand information processing networks (Glasser et al., 2016; Jain et al., 1998; Jain et al., 1994; Kuehn et al., 2017; Preuss et al., 1997; Van Essen et al., 2012). More recently, functional connectivity, particularly the correlation of intrinsic blood oxygenation level dependent (BOLD) fMRI signals from different brain areas, which is considered to represent functional association between regions, has been utilized for parcellation of the cortex (Gordon et al., 2016).

Resting-state functional connectivity is determined from correlations in the low frequency (< 0.01 Hz) fluctuations of BOLD fMRI signals between different regions in a resting brain revealing functionally related areas (Biswal et al., 1995). Such time series cross-correlation analysis between a seed region of interest (ROI) and voxels in rest of the brain reveals functional connections across different brain regions even without any explicit task paradigm (Biswal et al., 1995). The correlated sets of voxels give rise to functional ‘maps’, also referred to as functional connectivity networks. These have been found to be consistent within and between subjects (Choe et al., 2015; Laumann et al., 2015; Shehzad et al., 2009). The resting-state functional connectivity has been studied in a variety of species ranging from mice, ferrets, cats, and monkeys to humans (Hutchison et al., 2013; Stafford et al., 2014; Zhou et al., 2016). Seed-based resting-state fMRI (rsfMRI) analysis enables tailored investigation to delineate networks spanning the entire brain. Furthermore, subject specific ROI analysis using rsfMRI data can reveal regional differences within a cortical area, as well as individual variations across different subjects (Braga and Buckner, 2017).

In functional connectivity studies traditionally somatosensory and motor cortical areas have been lumped together (Beckmann et al., 2005; Hutchison et al., 2011; Mantini et al., 2013). This network includes different topographic representations in the primary somatosensory, motor and secondary somatosensory areas (Disbrow et al., 2000; Manger et al., 1996; Nelson et al., 1980; Stepniewska et al., 1993). However, it is known from anatomical connectivity studies that different body part representations form independent information processing modules. For example, the hand and face representations in the somatosensory cortex have different interconnectivity, sources of feedback connections and inter-hemispherical connections (Chand and Jain, 2015; Fang et al., 2002; Killackey et al., 1983; Liao et al., 2013). Callosal connectivity also varies widely for different body part representations, for example, the face and trunk representations have higher connectivity as compared to the hand and foot regions (Killackey et al., 1983).

Resting-state connectivity largely reflects direct anatomical connectivity between different brain regions (Damoiseaux and Greicius, 2009; van den Heuvel et al., 2009; Wang et al., 2013), as well as functional connectivity between areas not directly connected anatomically (Adachi et al., 2012; Vincent et al., 2007). Recent neuroimaging studies have considered differences in cortical function and functional connectivity to delineate different brain areas (Glasser et al., 2016; Gordon et al., 2016; Sohn et al., 2012). Functional connectivity studies have found topography dependent network subdivisions within the somatomotor network in humans (Power et al., 2011; Yeo et al., 2011). Utilizing *in vivo* cortical myelin mapping and resting-state analysis, Kuehn et al. (2017) showed that the functional connectivity patterns in humans also follow the architectonic differences reiterating the importance of body part representations as an organizing principle for functional networks.

We hypothesized that different body part representations within the primary somatosensory area 3b have different connectivity networks with different functionally correlating nodes. In order to test this hypothesis, we analyzed seed-based resting-state fMRI connectivity of different body part representations. Further, to determine if the correlated features of these networks are evolutionarily conserved, we compared functional connectivity profiles in humans and macaque monkeys.

## 2. METHODS

### 2.1. Subjects

Five adult male macaque monkeys (*Macaca mulatta*), 8-11 years of age and weighing between 8-10 kg were used. All animal procedures were approved by the Institutional Animal Ethics Committee of National Brain Research Centre, and the Committee for the Purpose of Control and Supervision of Experiments on Animals (CPCSEA), Government of India, and conformed to NIH guidelines for care and use of animals in biomedical research. Twenty three right-handed human subjects (10 females and 13 males) between the ages of 22-39 years took part in the study. Participants had no history of neurological or psychiatric illness (self-reported). Informed consent was obtained from all the human subjects. Human study protocols were approved by the Institutional Human Ethics Committee.

### 2.2. Data acquisition

#### 2.2.1 Macaque monkeys

For magnetic resonance data acquisition, macaque monkeys were initially anesthetized with ketamine hydrochloride (8 mg/kg, IM). Glycopyrrolate (0.015 mg/kg, IM) was administered in order to reduce the salivary secretions. When the monkeys were areflexive, they were intubated with an appropriately sized endotracheal tube and the anesthesia was switched to isoflurane (1-2% in oxygen, Surgivet CDS 2000). T1 weighted anatomical scans were acquired using 1-2% isoflurane in oxygen while the resting-state scans were taken using a lower isoflurane percentage (0.5-0.8% in oxygen).

MR scans were acquired by transmitting radio-frequency pulses from a quadrature body-coil inside a 3-T clinical MRI scanner (Philips Achieva, Netherlands), and receiving signals using an eight-channel phased array knee coil (MRI Device Corporation, WI, USA) and sensitivity-encoding (SENSE) parallel acquisition (Pruessmann et al., 1999). Human knee coil was used for monkeys due to a better filling factor, which improved signal-to-noise ratio and the image quality (Dutta et al., 2014). Anesthetized animals were placed in supine position inside the scanner with the head completely inside the receiver coil. Head movements inside the coil were minimized by padding the gaps between the head and the coil with polyethylene foam blocks. T1 weighted anatomical scans of the brain were acquired using a 3D multishot Turbo Field Echo sequence (TR = 8.8 ms; TE = 4.4 ms; flip angle = 8°; 0.5 mm x 0.5 mm x 0.5 mm resolution; 250 mm x 210 mm x 56 mm field of view; 500 x 360 matrix; 112 transverse slices). During the scans, physiological condition of the monkeys was continuously monitored with a MRI-compatible pulse-oximeter (Nonin 8600FO, USA), keeping constant isoflurane (0.5-0.8%) and the oxygen flow rate (1.5 l/min). On each scanning day, functional data acquisition was always preceded by reference and anatomical scans. This enabled similar time lag between induction of anesthesia and the start of the functional scans across different days of acquisition.

##### 2.2.1.1 Resting-state scans

Resting-state fMRI scans were acquired in oblique horizontal slice orientations, using a single-shot gradient echo echo-planar imaging (TR = 2800 ms; TE = 30 ms; flip angle = 90°; 1.5 mm x 1.5 mm in-plane resolution; 2 mm thick slices; 96 mm x 102 mm x 54 mm field of view; 64 x 64 matrix; number of excitations (NEX) = 2. Each resting-state scan session consisted of 140 functional volumes acquired over approximately 13 minutes.

From the five monkeys, a total of 32 scans were acquired. The number of sessions for different monkeys were 11, 8, 6, 4 and 3, which were acquired on different days.

##### 2.2.1.2. Functional localiser scans

For the functional location studies, the scanning parameters were similar to an earlier study from our lab (Dutta et al., 2014). Macaque monkeys were initially anesthetized using ketamine (8 mg/kg IM) and xylazine (0.4 mg/kg IM). Subsequent maintenance of anaesthetic depth as required during scanning was achieved using low doses of ketamine (1.5 mg/kg IM). Stimulation of the glabrous skin of the digits and palmar surface was done manually using a polyester brush at a frequency of approximately 2 Hz. Cutaneous stimulation to chin was delivered using a smaller brush at the same frequency (see Dutta et al., 2014, Jain et al 1997, 2008). The stimulation frequency was paced by steady counting with respect to a clock, and was always done by the same experimenter. The functional MRI scans were acquired using the same MRI scanner (3T, Philips Achieva, Netherlands) with oblique horizontal slice orientations, using a single-shot gradient echo echo-planar imaging (TR = 2800 ms; TE = 30 ms; flip angle = 90°; 1.5 mm x 1.5 mm in-plane resolution; 2 mm thick slices; NEX = 2; 96 mm x 102 mm x 54 mm field of view; 64 x 64 matrix). Acquisition paradigm using standard block design consisted of alternating rest (28s) and stimulation (28s) blocks. During the scans physiological condition of the monkeys was continuously monitored with a MRI-compatible pulse-oximeter (Nonin 8600FO, USA). In each session 140 functional volumes were acquired.

#### 2.2.2 Humans

Data from human subjects were acquired in the same 3-T MRI scanner but using an 8 channel SENSE head coil. Anatomical T1 weighted images were acquired for each subject using 3D multishot TFE sequence (TR = 8.4 ms; flip angle = 8°; FOV= 250 mm x 230 mm x150 mm; 252 x 211 matrix; 150 slices) before the functional scanning sessions.

##### 2.2.2.1 Resting-state scans

Resting-state functional scans were acquired with a T2* weighted gradient echo EPI sequence (TR = 2000 ms; TE = 30 ms; flip angle = 70°; 3 mm x 3 mm in-plane resolution; slice thickness 4 mm; FOV = 230 mm x 242 mm x 132 mm; 33 slices). Two hundred functional volumes were acquired during which the participants lay in supine position with palms facing downwards. They were instructed not to intentionally move any body part, keep their eyes closed and try not to indulge in any active thought process. Data was acquired in two successive sessions on a single day from each subject.

Of the 23 subjects, the data from two subjects (both females) were discarded due to excessive head motion and self-reported sleepiness during the resting scans. Data from additional four subjects (2 females, 2 males) were not considered due to involuntary hand movements during the acquisition. Data acquired in 34 scans from the remaining 17 subjects (6 females, 11 males) were used for further analyses.

##### 2.2.2.2. Functional localiser scans

For acquiring BOLD signals in response to peripheral stimulation in humans, the chin and the glabrous skin of the digits and palmar surface of the hand were stimulated manually using a polyester brush at a frequency of 2 Hz as for the monkey stimulation. The functional MRI scans were acquired using a single-shot gradient echo echo-planar imaging (TR = 2000 ms; TE = 30 ms; flip angle = 70°; 3 mm x 3 mm in plane resolution; slice thickness 4 mm; FOV = 230 mm x 242 mm x 132 mm; 33 slices). Two hundred volumes were acquired per session using standard block design consisting of alternating rest and stimulation blocks. The functional scans were not acquired on the day of the resting-state scans.

### 2.3. Pre-processing

Monkey and human data were processed using statistical parametric mapping (SPM8) software (http://www.fil.ion.ucl.ac.uk/spm/software/spm8) operating in Matlab 2013a platform (MathWorks Inc., MA, USA). The acquired structural images were aligned in the anterior commissure-posterior commissure (AC-PC) plane with coordinates of AC set to zero. After removal of the dummy scans, the functional images were slice time corrected to remove time lag between slices within a volume. These slice-time corrected functional volumes were head motion corrected using a six-parameter affine ‘rigid body’ transformation to minimize differences between each successive scan and the reference scan (the first scan in the time series). Motion corrected functional volumes of monkeys were co-registered with the corresponding high-resolution subject specific structural images. For humans, structural images were normalized to the standard template (ICBM 2009a Nonlinear Symmetric template; Fonov et al., 2009). Images were then visually inspected to check the registration.

For seed to voxel analysis (see below) data were smoothed with a 2 mm (macaques) or 8 mm (humans) Full Width at Half Maximum Gaussian kernels. For ROI-ROI analysis unsmoothed data was used to avoid possible spill over from the neighboring voxels due to large smoothing kernels.

Functional connectivity for both macaques and humans was determined using CONN toolbox (version 15.e) for SPM (Whitfield-Gabrieli and Nieto-Castanon, 2012). Nuisance covariates of cerebrospinal fluid and white-matter signals were modeled and removed following CompCor strategy (Behzadi et al., 2007), as implemented in CONN. Linear regression was performed where signals from the white matter and the cerebrospinal fluid, along with the global signal and the motion parameters were taken as covariates of no interest. This was followed by a temporal band pass filtering (0.008-0.09 Hz) to reduce low frequency drifts and the high frequency physiological noise. Before calculating bivariate correlation coefficients across brain regions, the processed data were despiked with hyperbolic tangent squash function and linearly detrended to remove low drift scanner noise.

For analysis of the activation fMRI data in monkeys, subject specific analysis was done as described before (Dutta et al., 2014). The stimulation epoch was represented using a box car model which was convolved with a haemodynamic response function (hrf) as implemented in SPM. For humans, the analysis was done in ICBM template space using normalized functional images. Motion estimates were included in the regression model as covariates of no interest. General linear model based univariate analysis yielded statistical parametric maps which were thresholded using uncorrected *p* < 0.005 for monkeys as reported earlier (Dutta et al., 2014). The cluster selection criterion was set to two or more contiguous voxels (see Dutta et al., 2014). In humans the activation clusters were thresholded using family wise error correction (FWER) at *p* < 0.05. Statistical maps were overlaid on the standard template images after transforming to the INIA19 standard space in monkeys and ICBM template in humans for comparative visualization (see Fig. 2).

### 2.4. Regions of Interest (ROIs)

For determining resting-state functional connectivity of different somatosensory representations, we demarcated ROI’s for area 3b, and the face representation (face3b), hand representation (hand3b), and rest of the medial region (med3b) in area 3b (Fig. 1). Other regions of interest (ROIs) for both monkeys (MK) and humans (HU) were demarcated as described below. The ROI’s were drawn on successive parasagittal slices and confirmed by reexamining the ROI’s in coronal and axial planes (Fig. 3).

**Fig. 1.**
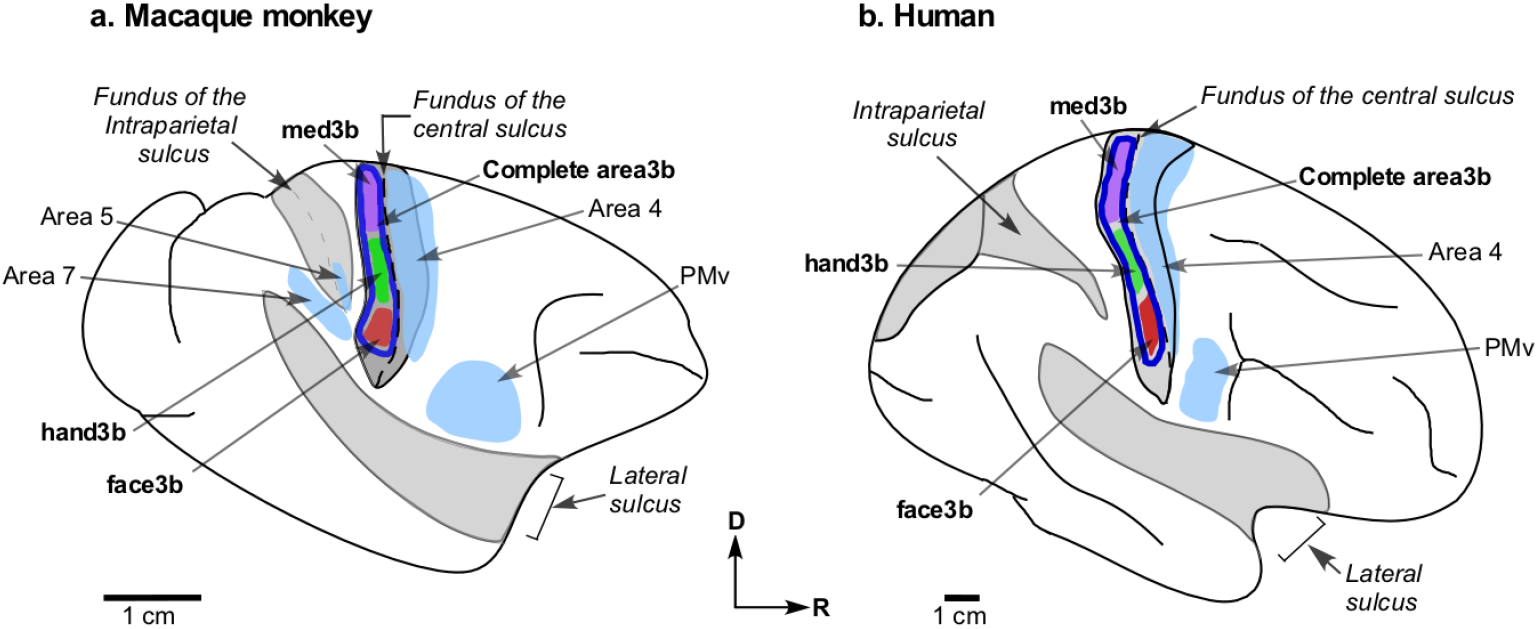
Schematic representation of the regions of interest (ROI’s) used for the seed-based resting-state functional connectivity analysis of the somatosensory cortex in (a) macaque monkeys and (b) humans. Complete area 3b ROI (thick dark blue outline; area3b), and ROIs of different body part representations - face3b (red), hand3b (green) and med3b (violet) are shown. The target ROI’s are light blue. The ROI’s are shown on outline drawings of lateral view of the cortical surface. The major sulci are labelled in italics for reference. Some of the sulci are shown opened (grey shading). S2 ROI is not visible in this view. PMv, Premotor Ventral Area; D, dorsal; R, rostral.

#### 2.4.1 Macaque monkeys

The ROI were drawn (Fig. 1a) on subject specific high resolution T1 images in both the hemispheres. The entire medio-lateral extent of the primary somatosensory area 3b on the anterior bank of the post central gyrus, excluding the medial wall representations was taken as the complete area 3b seed ROI (area3b). The ROI was restricted to grey matter using a grey matter mask. The boundaries between area 3b and 3a, and area 3b and 1 were demarcated with reference to the cytoarchitectonic and electrophysiology studies in monkeys (Chand and Jain, 2015; Jain et al., 2008; Nelson et al., 1980; Tandon et al., 2009). From these data area 3b was judged to lie between the depths of 1.5 mm-8 mm from the lip of the central sulcus (Fig. 3). To ensure that area 3b does not extend into the adjacent areas, voxels adjacent to the rostral boundary with area 3a and caudal boundary with area 1 were scrubbed off.

The ROI’s for body part representations (Fig. 3) were drawn within limits of area 3b boundaries with reference to the published electrophysiological and anatomical studies, and monkey atlases (Chand and Jain, 2015; Jain et al., 2008; Nelson et al., 1980; Paxinos et al., 2000; Saleem and Logothetis, 2006). A perpendicular from the tip of the intraparietal sulcus to the central sulcus on the surface of the brain was considered as the hand-face border, i.e. the medial limiting boundary of the face ROI (Chand and Jain, 2015; Jain et al., 2008; Manger et al., 1997). The face ROI included the contralateral representations of face regions and excluded the antero-lateral representations of ipsilateral trigeminal and intraoral inputs (Manger et al., 1996). The hand ROI was restricted to 7 mm distance medial from the hand-face border (Jain et al., 2008; Kambi et al., 2011, 2014; Nelson et al., 1980; Tandon et al, 2009). This included the hand representation i.e., digits and palm, but excluded other forelimb representations. Accuracy of the hand and chin ROI’s was further confirmed by referencing the fMRI activation loci in response to the chin and the hand stimulation in the same monkeys (Fig. 2a; also see Dutta et al., 2014).

**Fig. 2.**
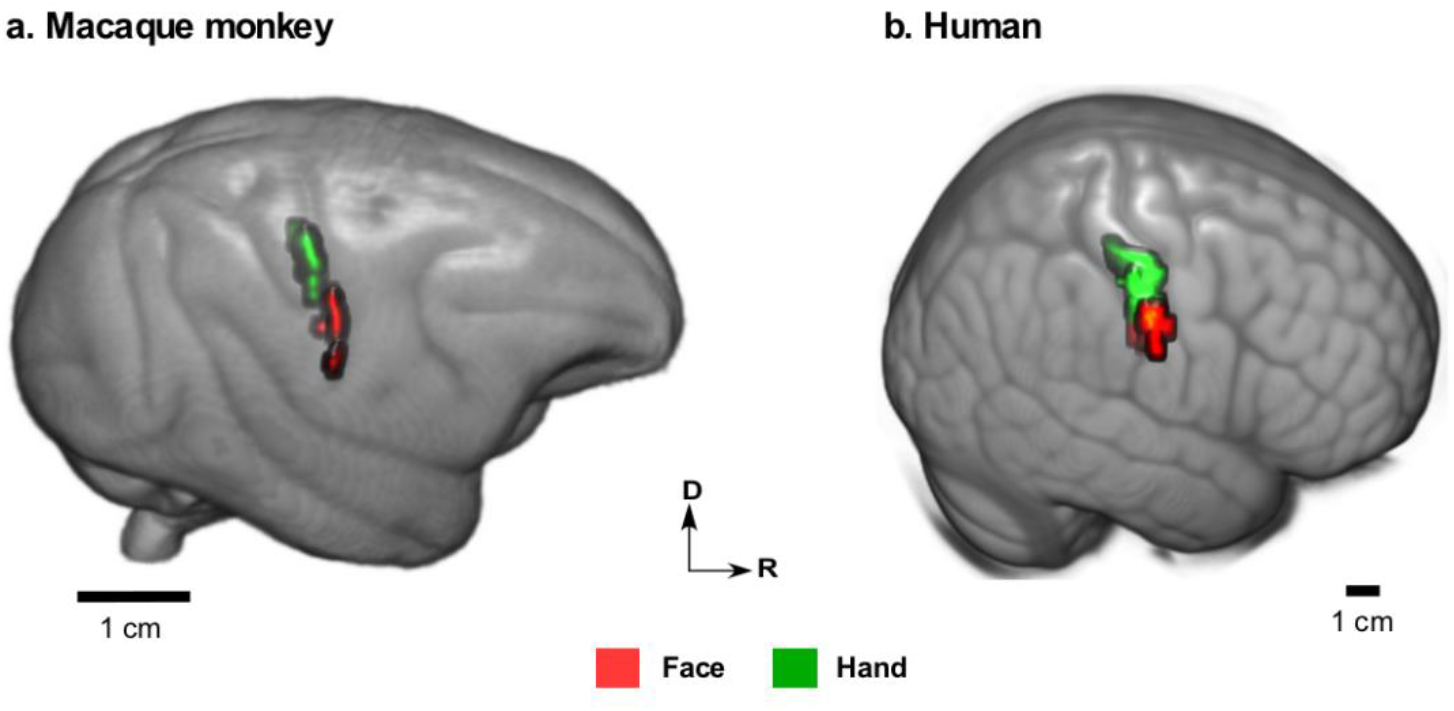
Voxels activated in fMRI scans of **(a)** macaque monkeys (n = 5) and **(b)** humans (n = 5) when the hand, i.e. glabrous digits and palm (green) or the face (red) was undergoing tactile stimulation. The voxels with peak activation in the post central gyrus are rendered on respective standard brain template image. All other brain regions are masked out. Statistical thresholds are *p* < 0.005 (uncorrected) for monkeys and *p* < 0.05 (FWER corrected) for humans. D, dorsal; R, rostral.

The remaining medial region of area 3b excluding the medial wall was considered together as the third ROI termed med3b. This included representations of the shoulder, trunk, foot and parts of the leg that are on the dorsal surface (Jain et al., 2008; Nelson et al., 1980). This ROI also included medial-most parts of the upper arm representation. Due to the uncertainties in demarcating boundaries between these representations, the combined ROI was used. Care was taken to avoid voxel overlap between face3b, hand3b, and med3b ROIs in the medio-lateral direction as well by having adequate gap between the representations by scrubbing off voxels.

Other ROIs, the primary motor cortex (area4), ventral premotor cortex (PMv) and secondary somatosensory cortex (S2) were drawn with reference to the published anatomical and electrophysiological data, and macaque brain atlases (Barbas and Pandya, 1987; Burton et al., 1995; Krubitzer et al., 1995; Paxinos et al., 2000; Saleem and Logothetis, 2006). ROIs were drawn for the body part representations in area 4 to demarcate the face (face4), the hand (hand4) and the remaining medial region (med4) in accordance with the published reports (Stepniewska et al., 1993).

#### 2.4.2 Humans

The ROIs were delineated on the standard brain template (Fonov et al., 2009). The complete area 3b ROI (area3b) included the deeply situated area 3b on the posterior bank of the central sulcus extending from the lateral face area to medial boundary where the central sulcus meets the midline sulcus (Blankenburg et al., 2003). This ROI excluded the medial wall representations of area 3b (Fig. 1b). Lateral boundary of area 3b extended upto the lateral boundary of the face ROI (see below). The voxels included in the area 3b ROI was confined within the grey matter. A grey matter mask was used to exclude white matter voxels. The boundaries between area 3b and area 1, area 3b and area 3a were delineated with reference to published anatomical studies, SPM Anatomy toolbox and human brain atlas (Bakker et al., 2015; Eickhoff et al., 2005; Geyer et al., 1999, 2000b). Based on these rostral boundary of area 3b was estimated to lie at depths varying from 7 mm from the lip of the central sulcus in the region of the trunk representation to 40 mm near the hand representation (Fig. 3).

**Fig. 3.**
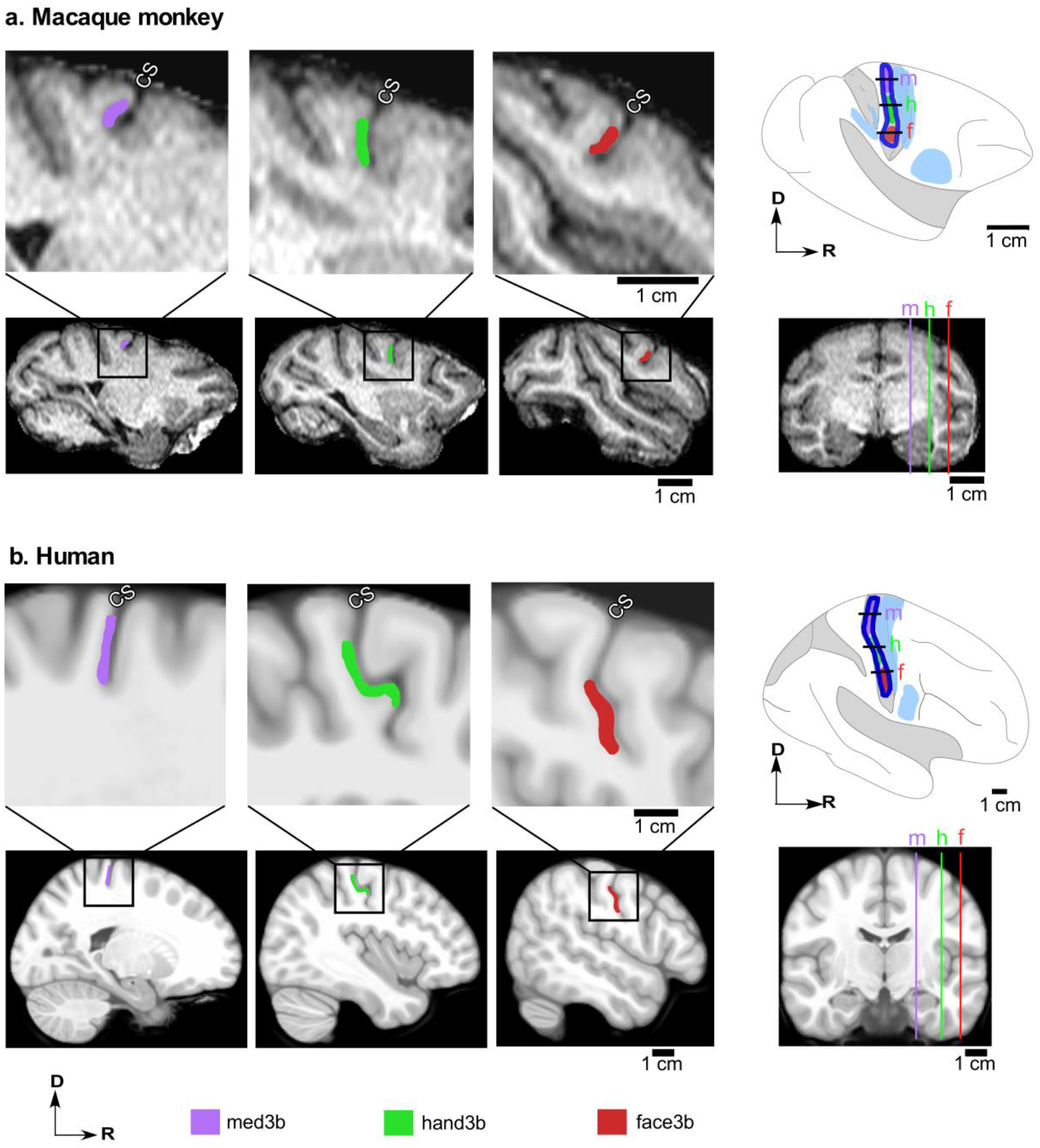
Examples showing how region of interests (ROI’s) used for functional connectivity analysis were drawn in area 3b of **(a)** macaque monkeys and **(b)** humans. The ROI’s were drawn on the subject specific high resolution T1 structural image in monkey and template brain in humans. Upper row on the left in ‘a’ and ‘b’ show ROI’s for the face (red), hand (green) and medial representations (violet) marked on representative sagittal slices shown enlarged in the region of the central sulcus (CS). Second rows in ‘a’ and ‘b’ on the left show complete sagittal slices with boxes marking the region shown enlarged in the upper rows. Upper right in ‘a’ and ‘b’ show locations of the slices marked by horizontal black lines on drawings of the dorsolateral surface of the brains to show their position with respect to the CS. Locations of slices are also shown as color coded lines on the coronal slices of the brains in the lower right (f, face3b; h, hand3b; m, med3b). D, dorsal; R, rostral.

The hand (hand3b) and the face (face3b) ROIs were then delineated. The anterior and posterior extents of the face3b and hand3b were drawn according to the published anatomical studies (Geyer et al., 1999, 2000b). The inverted Ω shaped knob was taken as suggestive of the motor hand area (Yousry et al., 1997). The somatosensory hand representation presumed to be lying in apposition to the area 4 hand region was drawn on the post central gyrus with the ‘knob’ as a guide. Other imaging studies were also used as a reference for location of the hand area (Blankenburg et al., 2003; Maldjian et al., 1999; Nakamura et al., 1998). Face ROI was drawn lateral to the hand ROI taking care to exclude the very lateral representations of intra-oral regions and tongue (Miyamoto et al., 2006). The lateral part of the face representation overlies the deeper medial part of the tongue representation, which was carefully excluded. We acquired fMRI data while stimulating the entire hand (i.e. glabrous digits and palm) or the chin, and used these data to confirm the placement of the ROI’s (Fig. 2b).

Similar to monkeys, med3b in humans comprised of parts of area 3b medial to the hand representation up to the location where central sulcus meets the midline, and excluded the medial wall representations. Voxels near the ROI edges in area 3b were scrubbed off to avoid mediolateral overlap between them.

Other ROIs - area4, PMv and S2 were also drawn on the standard ICBM template. ROI’s for area 4 (the primary motor cortex) and PMv (ventral premotor cortex) were drawn using SPM Anatomy toolbox with reference to the available anatomical data (Eickhoff et al., 2005; Geyer et al., 1996, 2000a, 2004). Secondary somatosensory ROI, which included both S2 and PV, was drawn using SPM Anatomy toolbox and referring to the published literature on S2 topography (Blatow et al., 2007; Disbrow et al., 2000; Eickhoff et al., 2005, 2007; Ferretti et al., 2004).

ROIs were drawn for the body part sub-divisions of the primary motor cortex to demarcate the face (face4), the hand (hand4) and the remaining medial region (med4) in accordance with the published reports (Geyer et al., 2000a; Schieber, 2001).

### 2.5. Connectivity analysis

For both monkeys and humans, seed-to-voxel connectivity was determined using CONN toolbox. For analysis in each human subject, the two successive sessions, which were acquired on the same day were concatenated. We did preliminary analysis taking the first and the second sessions of all the subjects separately. However, since no significant differences was found between the two sessions (not shown), the sessions were concatenated. Each session of the monkeys, which were all acquired on separate days, was analyzed separately. Data from each ROI were averaged across the sessions for each animal. To determine connectivity maps complete area 3b, face3b, hand3b and med3b ROI’s were used as seed regions. While performing linear regression in CONN, motion parameters were taken as regressors. Resultant *beta* maps, which were Fisher-z transformed correlation value maps, were used for further statistical analysis.

#### 2.5.1 Seed-to-voxel analysis

To perform group analysis for monkeys, the subject level *beta* maps were transformed to standard INIA19 template space (Rohlfing et al., 2012) using transformation matrix. The transformation matrix was generated using FSL’s linear and non-linear registration tool FLIRT and FNIRT respectively, registering subject specific high resolution T1 image to INIA19 template space (Jenkinson et al., 2002). Transformed *beta* maps in the template space were statistically tested using one sample t-test with null hypothesis of no correlation at *p* < 0.01 with False Discovery Rate (FDR) correction. Resultant group level statistical maps were then mapped on inflated brain surface using CARET (ver. 5.616; Van Essen et al., 2001) and were displayed on slices using MRIcron software (ver. 1; Rorden and Brett, 2000).

For human subjects, subject level *beta* maps were used for the statistical group analysis performed using one sample t-test with null hypothesis of no correlation thresholded at *p* < 0.01 with FDR correction. The group level statistical maps were mapped on inflated standard template brain surface using BrainNet Viewer (Xia et al., 2013) and were displayed on slices using MRIcron software (ver. 1; Rorden and Brett, 2000).

#### 2.5.2 ROI-ROI analysis

To calculate the effect size of highly correlating ROIs, specific ROI-ROI Fisher-z transformed correlations were calculated between different ROI’s using CONN. One sample t-test was performed at the subject level in both monkeys and humans with null hypothesis of no correlation with a threshold for significance set for monkeys at *p* < 0.05, and humans at *p* < 0.00001. The thresholds were decided empirically referring to the connectivity networks reported for the somatomotor cortex (Biswal et al., 1995; Kuehn et al., 2017; Mantini et al., 2013; Vincent et al., 2007; see ‘limitations of the study’ in ‘Discussion’). Goal was to avoid spurious connectivity while ensuring that the known connectivity patterns were not thresholded out.

To statistically compare the connectivity of ROI-ROI pairs, two-way ANOVA (alpha = 0.05) and post hoc Tukey test (*p* < 0.01) was implemented using GraphPad Prism 6.01 for Windows (GraphPad Software, California USA, www.graphpad.com). All other ROI-ROI statistical tests were performed in MATLAB.

## 3. RESULTS

We determined resting-state functional connectivity networks of different topographic representations in area 3b of macaque monkeys and humans. Exploratory seed-to-voxel correlation analysis was first done to reveal functionally connected regions in the brain. The networks thus revealed were further analyzed by ROI-ROI correlation analysis. We describe below our results for macaque monkeys and humans together for ease of comparison.

### 3.1. Somatosensory resting-state networks: seed-to-voxel correlation analysis

Resting-state network organization was determined taking the entire area 3b or different body part representations as seeds for the exploratory correlation analysis. The networks were broadly similar for both humans and monkeys (Fig. 4 and 5).

**Fig. 4.**
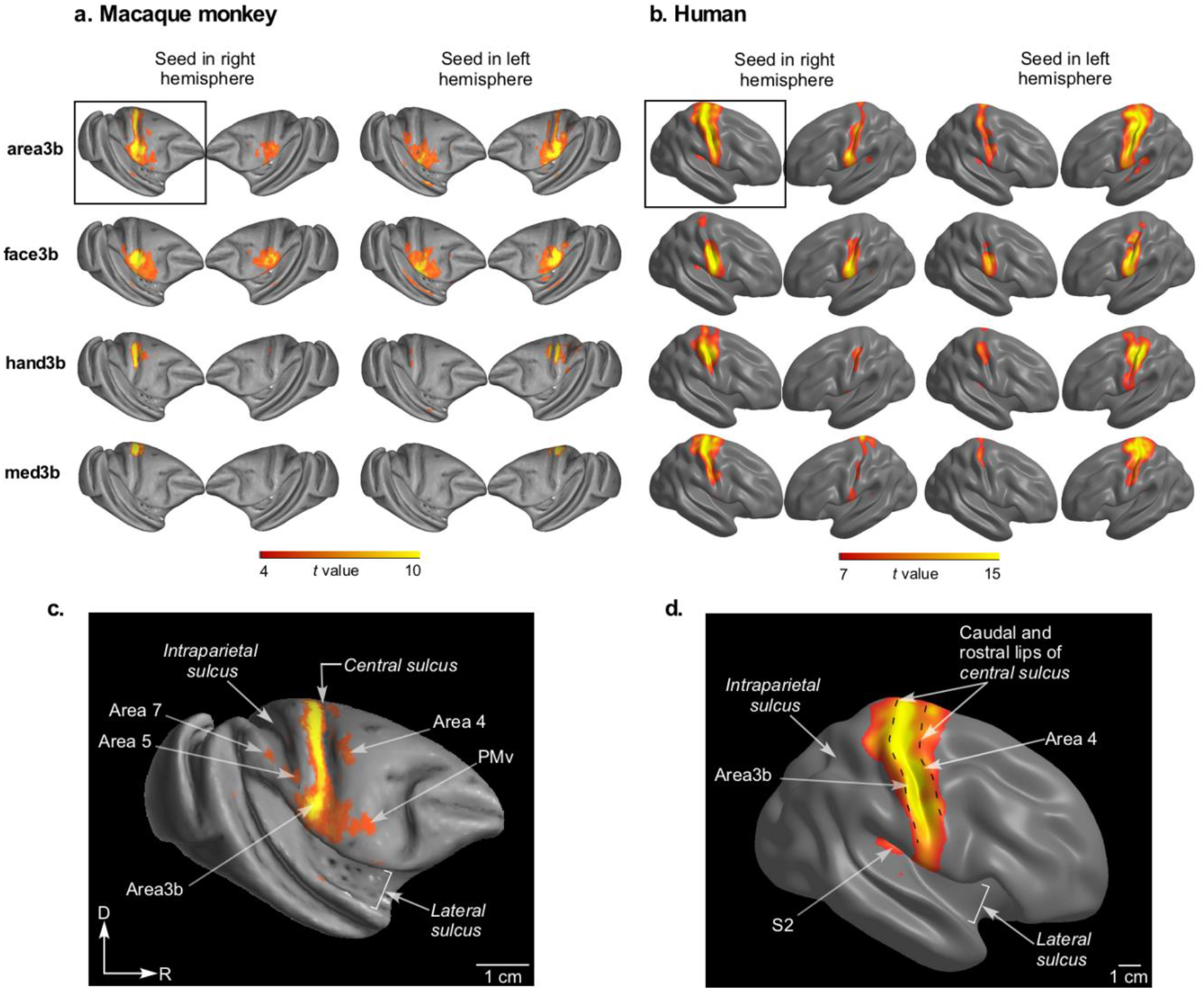
Seed to voxel resting-state functional connectivity of the somatosensory cortex of (**a, c**) macaque monkeys and (**b, d**) humans. (**a** and **b**) Bi-hemispherical views of the somatosensory network in macaques and humans shown on partially inflated cortical surfaces when the seed was complete area 3b (area3b; top row), face representation in area 3b (face3b; second row), the hand representation (hand3b; third row), or the medial part of area 3b (med3b; bottom row). To illustrate symmetrical nature of the network, data are shown for seed in the left as well as the right hemisphere (see labels on top). The ‘t-values’ correspond to statistical significance of *p* < 0.01 (FDR corrected). Note that t-values are scaled to highlight the differences in connectivity maps of ROI’s (see the colour scale bars at the bottom).’**c’** and ‘**d**’ show enlarged view of the boxed figurines in ‘a’ and ‘b’ respectively to illustrate the details. D, dorsal; R, rostral; orientation arrows shown in ‘c’ also applies to ‘d’.

**Fig. 5.**
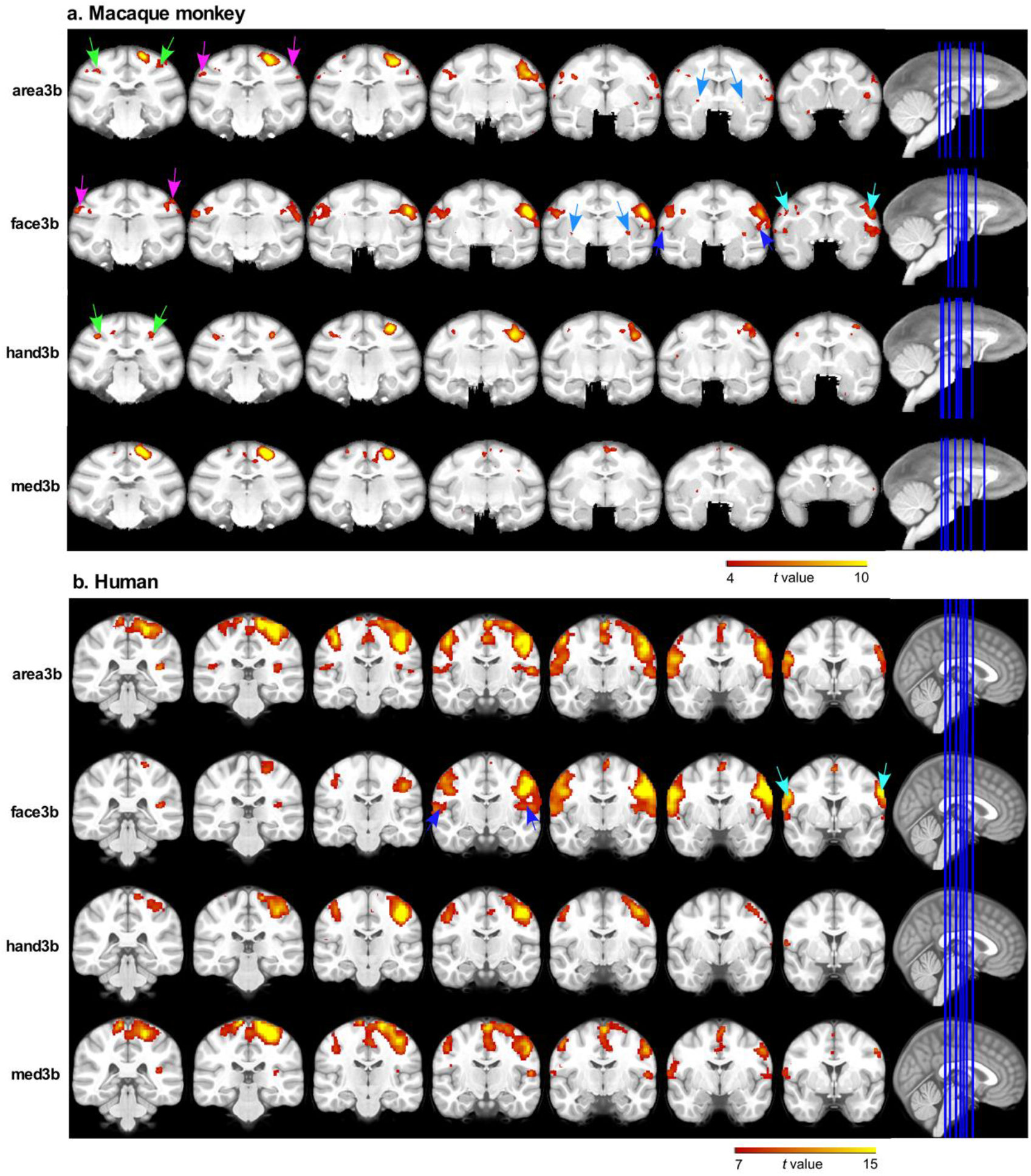
Seed to voxel correlations in area 3b of the right hemisphere of (**a**) monkeys and (**b**) humans shown on a series of coronal slices. The slices are arranged in the caudal to rostral order from left to right. The right-most column shows parasagittal sections of the brain with the blue lines marking the plane from which the coronal slices are taken. Correlations with area 7 (pink arrows), area 5 (green arrows) and putamen (blue arrows) were observed only in monkeys. Locations of S2 (violet arrows) and PMv (cyan arrows) are also marked. M, medial; R, rostral. Other conventions as for Figure 4.

The network of complete area 3b (area3b) in both the primate species comprised of contralateral area 3b. In both the hemispheres area3b network also had nodes in area 4, second somatosensory area (S2), premotor ventral area (PMv) and insula (Fig. 5). In monkeys it also included area 7, putamen and area 5 of both the hemispheres (Fig. 5a).

Further analysis of connectivity of different body part representations revealed that each representation had a distinct connectivity pattern as described below. The face representation (face3b), in both monkeys and humans showed strong connectivity with the face representation in contralateral area 3b. Face3b also showed bilateral connectivity with the face representation in area 4 (face4), PMv, S2, and insula (Fig. 4 and 5). However, only in monkeys, face3b showed bilateral connectivity with area 7 and putamen (Fig. 5a).

The other two ROIs i.e. hand3b and med3b revealed a network with fewer nodes than face3b. In monkeys the hand3b network included area 5 and the hand representation in area 4 (hand4) of both the hemispheres. The human hand3b showed connectivity to the contralateral hand3b and the ipsilateral hand4 (Fig. 4 and 5). Med3b in both monkeys and humans showed connections to the rostrally adjacent medial part of area 4 (med4) and the contralateral med3b (Fig. 5).

The results showed that the functional connectivity network of area 3b is different for different body part representations. Furthermore, a seed ROI placed in either hemisphere had generally similar connectivity pattern, suggesting that the networks are largely laterality independent (Fig. 4).

The nodes of the network revealed by seed-to-voxel connectivity results guided the ROI-ROI analysis described below.

### 3.2. ROI-ROI correlations: Different body part representations in area 3b contribute differentially to the complete area 3b functional connectivity

For a detailed investigation of the results of the seed-to-voxel analysis, ROI-ROI analysis was performed. ROIs were drawn in the seed as well as the target areas in the primary somatosensory and motor areas, S2 and PMv. In order to determine the extent of connectivity between correlated regions BOLD signals from all the voxels in each ROI were averaged and time series correlations of the averaged signals were determined for each ROI-ROI pair. Color coded connectivity matrices were constructed using averaged Fisher-z transformed correlation coefficients (*CC_7_*) for these ROI pairs in both humans and macaque monkeys to illustrate the results (Fig. 6).

**Fig. 6.**
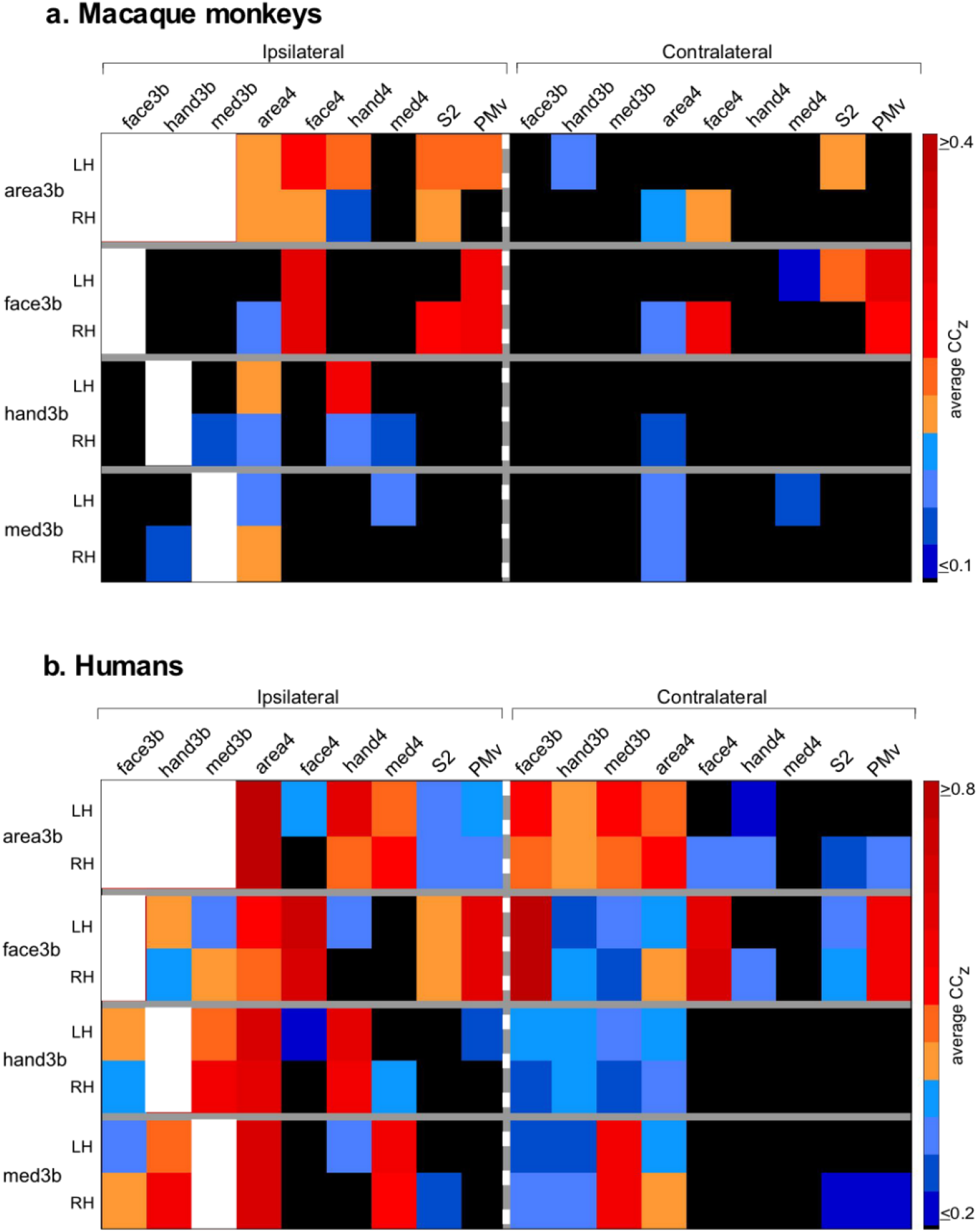
Correlation matrices showing ROI-ROI resting state functional connectivity between different ROI’s in ipsilateral (left of the dashed white line) and contralateral hemispheres (right of the dashed white line) of (a) monkeys and (b) humans. The seed regions are shown on the y-axis, and the target regions on top. Each colour coded value denotes averaged Fisher-z transformed correlations (CCz) acquired from different subjects. For colour codes, see the bars on the right. Statistically nonsignificant correlations are shown in black and autocorrelations are in white (*p* ≤ 0.05 for monkeys; *p* ≤ 0.00001 for humans; one sample t-test). 3b, area 3b; 4, area 4; ‘face’, ‘hand’, ‘med’ prefixes denote the face, hand and the medial part of area 3b or 4. LH, left hemisphere; RH, right hemisphere.

In both the species ROI encompassing entire area 3b in either of the hemisphere revealed significant connections with the contralateral area 3b. Area 3b in the right hemisphere also had significant bilateral correlations with area 4 (Fig. 6). In humans, area 3b of right hemisphere also showed bilateral connectivity with S2 and PMv, whereas area 3b of the left hemisphere has connectivity with only ipsilateral S2 and PMv. Area 3b of the left hemisphere in monkeys showed correlations with bilateral S2 and ipsilateral area 4 (Fig. 6).

Looking at the connectivity of individual body part representations, in both the species, face3b in the right hemisphere showed significant bilateral connectivity with face representation in area 4 and PMv. In monkeys face3b of the right hemisphere had significant connectivity with ipsilateral S2, and face3b of the left hemisphere with contralateral S2. This essentially means that right S2 had connectivity with face3b in both the hemispheres. In humans, face3b had significant connections with the contralateral face3b, and bilateral connectivity with S2 (Fig. 6).

The hand3b in both species had significant connectivity with ipsilateral hand4. Med3b in both hemispheres of humans and the left hemisphere of monkeys had significant correlations to homotopic representation in area 4 (one sample t-test, for *p*-values see Tables 1 and 2).

**Table 1.**
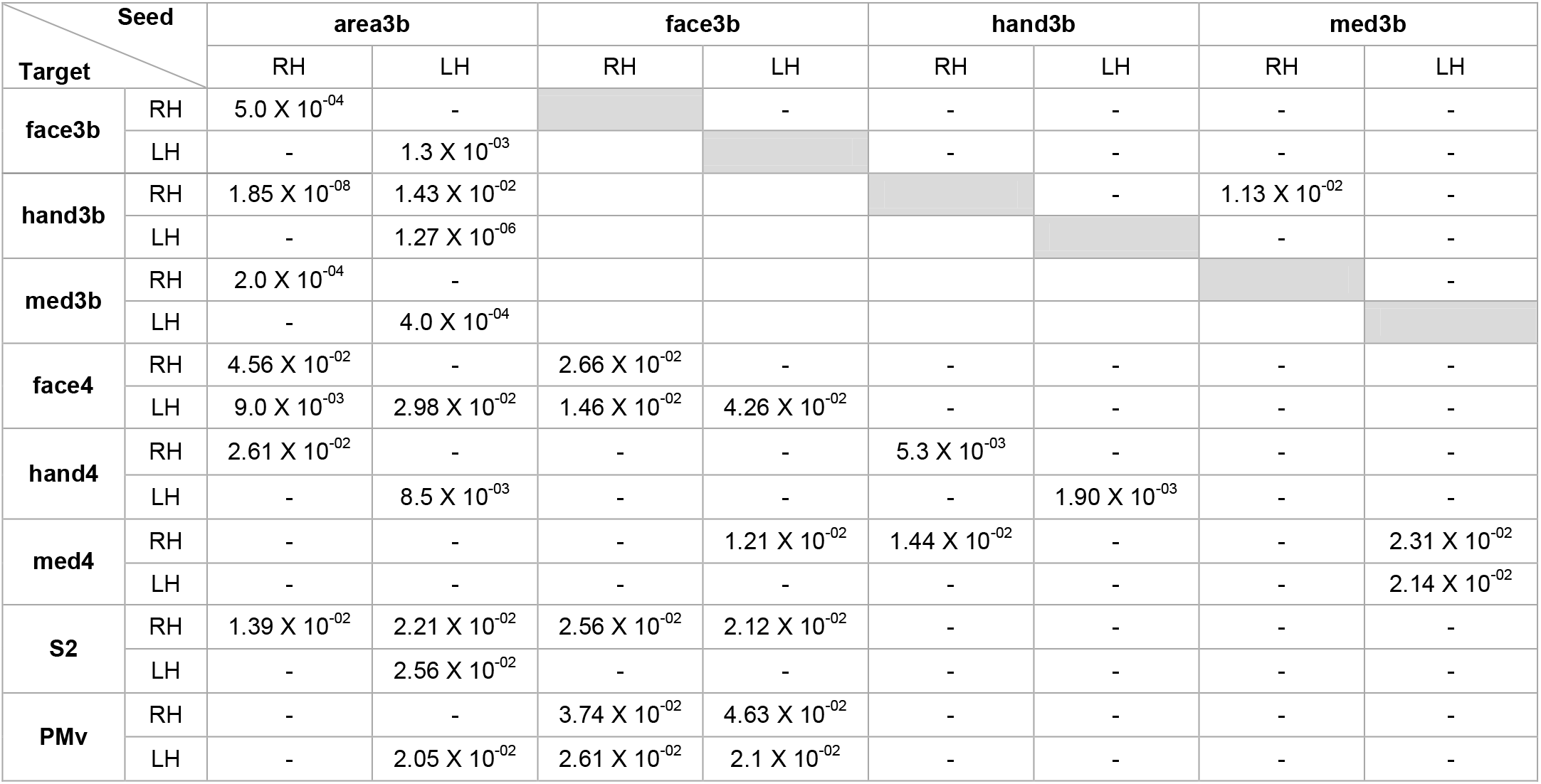
*p*-values for ROI-ROI correlations for monkeys. LH, left hemisphere; RH, right hemisphere. Only significant pair-wise correlations i.e. where *p-value* < 0.05 (one sample t-test) are shown. Grey boxes are auto-correlations.

**Table 2.** *p*-values for ROI-ROI correlations for humans. LH, left hemisphere; RH, right hemisphere. Only significant pair-wise correlations i.e. where *p-value* < 0.00001 (one sample t-test) are shown. Grey boxes are auto-correlations.

| Seed \ Target |  | area3b |  | face3b |  | hand3b |  | med3b |  |
| --- | --- | --- | --- | --- | --- | --- | --- | --- | --- |
|  |  | RH | LH | RH | LH | RH | LH | RH | LH |
| face3b | RH | $3.70 \times 10^{-12}$ | $3.13 \times 10^{-09}$ | | $6.66 \times 10^{-10}$ | $5.80 \times 10^{-08}$ | $3.80 \times 10^{-06}$ | $5.86 \times 10^{-09}$ | $6.21 \times 10^{-06}$ |
| | LH | $3.19 \times 10^{-08}$ | $5.85 \times 10^{-12}$ | | | $4.82 \times 10^{-06}$ | $1.91 \times 10^{-08}$ | $6.70 \times 10^{-06}$ | $1.71 \times 10^{-07}$ |
| hand3b | RH | $6.94 \times 10^{-14}$ | $1.91 \times 10^{-08}$ | | | | $4.03 \times 10^{-07}$ | $8.03 \times 10^{-10}$ | $3.51 \times 10^{-07}$ |
| | LH | $8.38 \times 10^{-09}$ | $1.92 \times 10^{-12}$ | | | | | $5.15 \times 10^{-06}$ | $8.35 \times 10^{-09}$ |
| med3b | RH | $5.40 \times 10^{-14}$ | $1.66 \times 10^{-09}$ | | | | | | $2.34 \times 10^{-09}$ |
| | LH | $1.00 \times 10^{-08}$ | $1.68 \times 10^{-15}$ | | | | | | |
| face4 | RH | - | - | $1.18 \times 10^{-08}$ | $2.53 \times 10^{-06}$ | - | - | - | - |
| | LH | $5.90 \times 10^{-08}$ | $1.92 \times 10^{-08}$ | $9.59 \times 10^{-09}$ | $1.29 \times 10^{-08}$ | - | $9.64 \times 10^{-06}$ | - | - |
| hand4 | RH | $2.84 \times 10^{-07}$ | $1.40 \times 10^{-06}$ | - | - | $1.07 \times 10^{-07}$ | - | - | - |
| | LH | $8.16 \times 10^{-06}$ | $1.68 \times 10^{-08}$ | $2.44 \times 10^{-06}$ | $2.72 \times 10^{-07}$ | - | $2.77 \times 10^{-08}$ | - | $1.35 \times 10^{-07}$ |
| med4 | RH | $3.39 \times 10^{-07}$ | - | - | - | $9.68 \times 10^{-07}$ | - | $2.52 \times 10^{-06}$ | - |
| | LH | - | $1.79 \times 10^{-07}$ | - | - | - | - | - | $3.56 \times 10^{-07}$ |
| S2 | RH | $2.54 \times 10^{-07}$ | - | $1.00 \times 10^{-08}$ | $4.87 \times 10^{-06}$ | - | - | $4.48 \times 10^{-06}$ | - |
| | LH | $5.04 \times 10^{-07}$ | $4.36 \times 10^{-06}$ | $5.86 \times 10^{-07}$ | $3.08 \times 10^{-06}$ | - | - | $8.91 \times 10^{-06}$ | - |
| PMv | RH | $9.49 \times 10^{-07}$ | - | $8.76 \times 10^{-09}$ | $1.73 \times 10^{-06}$ | - | - | - | - |
| | LH | $1.21 \times 10^{-07}$ | $2.52 \times 10^{-09}$ | $1.46 \times 10^{-08}$ | $4.71 \times 10^{-09}$ | - | $8.40 \times 10^{-06}$ | $8.94 \times 10^{-06}$ | - |

Thus ROI-ROI analysis revealed that face3b has more widespread and complex functional network than med3b and hand3b. The network of complete area 3b ROI largely includes networks of all the three representations viz. face3b, hand3b and med3b, although some of the nodes are seen only when individual body parts are considered separately. For example, in monkeys bilateral connectivity with PMv, and in humans bilateral connectivity with face4 is observed only when face3b is considered separately. Moreover, there are few nodes that show significant connectivity only when the entire 3b is considered to together (for details see Fig. 6).

Some of the salient features of the functional networks are described below.

#### 3.2.1 Connectivity with contralateral 3b

Different body part representations showed differences in the extent of connectivity with homotopic representations in the contralateral 3b. All ROI’s in humans showed significant bilateral connectivity (Fig. 6b). Average Fisher’s z-transformed correlation coefficient (CCz) values in humans showed that the highest inter-hemispherical correlation was for face3b, followed by med3b and hand3b (Fig. 7; face3b: *p* = 6.7 x 10^−10^; med3b: *p* = 2.3 x 10^−9^; hand3b: *p* = 4.03 x 10^−7^; one sample t-test). The ROI’s in monkeys although did not have statistically significant homotopic bilateral connectivity, but followed similar trend in the strength of connectivity (Fig. 7; face3b: *p* = 6.9 x 10^−2^; med3b: *p* = 1.08 x 10^−^; hand3b: *p* = 2.43 x 10^−1^; one sample t-test).

**Fig. 7.**
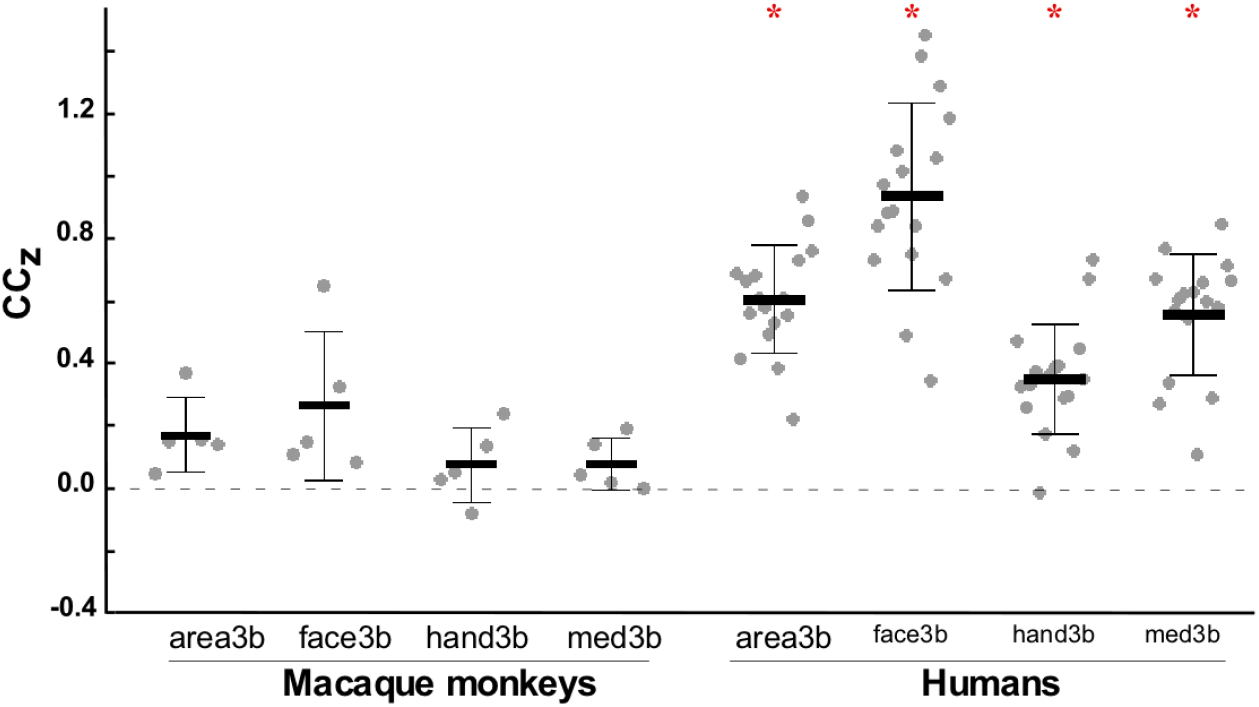
Inter-hemispheric homotopic functional connectivity of somatosensory area 3b, and ROI’s for different body part representations in macaque monkeys and humans. Each dot on the plot denotes Fisher-z transformed correlation value (CC_Z_) for a single subject in monkeys and humans. Thick horizontal lines shows the mean and the thin lines, ± SD. Asterisk* denotes significant statistical difference (*p* < 0.00001 in humans, one sample t-test).

The interhemispheric connectivity of all the three ROI’s - face3b, hand3b and med3b with non-homotopic ROI’s in contralateral area 3b was also significant only in humans (Fig. 6 and 10; one-sample t-test, see Table 2 for *p*-values).

#### 3.2.2 Functional connectivity with S2 and PMv

Our ROI’s in S2 and PMv encompassed the entire areas; no attempt was made to place ROI’s in representations of specific body parts. Therefore, the connectivity of individual body part representations that we refer to below is with the entire area S2 or PMv.

Face3b of humans in both hemispheres had significant connectivity with S2 bilaterally (see Fig. 8a for ipsilateral connectivity, LH, *p* = 3.08 x 10^−6^; RH, *p* = *p* = 1.0 x 10^−8^; for *p* values for contralateral S2 see Table 2). However, in monkeys face3b of the right hemisphere had significant connectivity only with ipsilateral S2 (Fig. 8a; *p* = 2.56 x 10^−2^), whereas the face3b of the left hemisphere had significant connectivity with only the contralateral S2 (Fig. 6; *p* = 2.12 x 10^−2^). Thus, S2 of only the right hemisphere was connected to face3b.

**Fig. 8.**
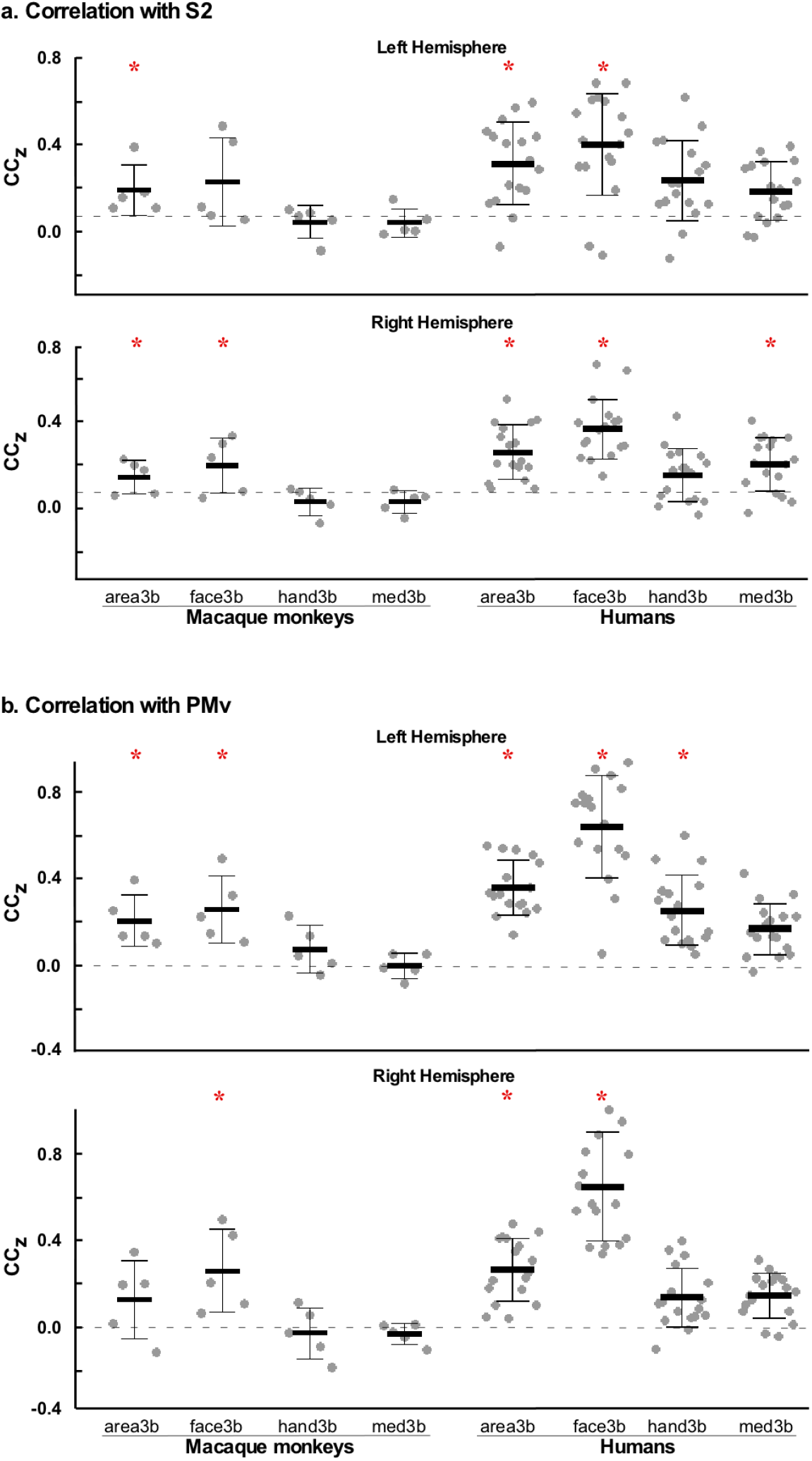
Correlation coefficients (CCz) of somatosensory ROIs with ipsilateral (a) S2 and (b) PMv in macaque monkeys and humans. Each dot on the plot denotes Fisher-z transformed correlation value (CCz) for a single subject in monkeys and humans. Horizontal lines show the mean (thick lines) and ± S.D (thin lines) of correlation coefficients. Asterisk* denotes statistical significant difference (*p* < 0.05 in monkeys; *p* < 0.00001 in humans, one sample t-test).

The hand3b of either monkeys or humans did not have significant connectivity with S2 (Fig. 8a; for monkeys LH, *p* = 3.15 x 10^−1^; RH, *p* = 4.18 x 10^−1^; for humans LH, *p* = 1.22 x 10^−4^; RH, *p* = 1.05 x 10^−4^; one sample t-test). Similar to hand3b, the med3b of monkeys showed no significant connectivity with S2 (Fig. 8a; LH, *p* = 3.0 x 10^−1^; RH, *p* = 3.05 x 10^−1^). However, in humans med3b of the right hemisphere had bilateral connectivity with S2, whereas med3b of the left hemispheres had no significant connectivity with S2 (Fig. 8a; LH, *p* = 3.89 x 10^−5^; RH, ipsilateral, *p* = 4.48 x 10^−6^, contralateral (see Fig 6b), *p* = 8.91 x 10^−6^; one sample t-test).

With PMv, face 3b of both monkeys and humans had significant connectivity (Fig. 8b; for monkeys LH, *p* = 2.1 x 10^−2^; RH, *p* = 3.74 x 10^−2^; for humans LH, *p* = 4.71 x 10^−9^; RH, *p* = 3=8.76 x 10^−9^; one sample t-test; for *p* values for contralateral PMv see Table 1 and 2). In monkeys the hand3b or med3b did not show any significant connectivity with PMv. In humans the hand3b of the left hemisphere had significant connectivity with ipsilateral PMv (Fig. 8b, top; *p* = 8.4 x 10^−6^), and med3b of only the right hemisphere had significant connectivity but only with contralateral PMv (Fig. 6b; *p* = 8.94 x 10^−6^).

The data suggested that in both the species the observed connectivity for the entire area 3b with S2 and PMv was primarily a reflection of the face3b correlations with these areas.

#### 3.2.3 Connectivity with homotopic representations in area 4

We compared the correlations of different body part representations in area 3b with the homotopic body part representation in motor area 4. All three ROIs - face3b, hand3b and med3b showed strong connectivity with the corresponding representation in the ipsilateral primary motor cortex in both macaque monkeys and humans, except for the right med3b of monkeys (Fig. 6). Interestingly, the mean correlation strength of face3b with face4 was significantly higher than the correlation of face3b with hand3b or med3b (see Fig. 9, left, for LH, macaque, *F*_(2, 81)_ = 25.84, *p* < 0.0001; humans, *F*_(2, 51)_ = 35.40, *p* < 0.0001; two-way ANOVA; *p* < 0.01, *post hoc* Tukey test; see Fig. 9, right, for RH, macaque, *F*_(2, 81)_ = 13.79, *p* < 0.0001; humans, *p* = *F*_(2, 51)_ = 30.56, *p* < 0.0001; two-way ANOVA; *p* < 0.01, *post hoc* Tukey test). Thus, connectivity of face representation in area 3b was stronger with face representation in area 4 than for other ROI’s in area 3b.

**Fig. 9.**
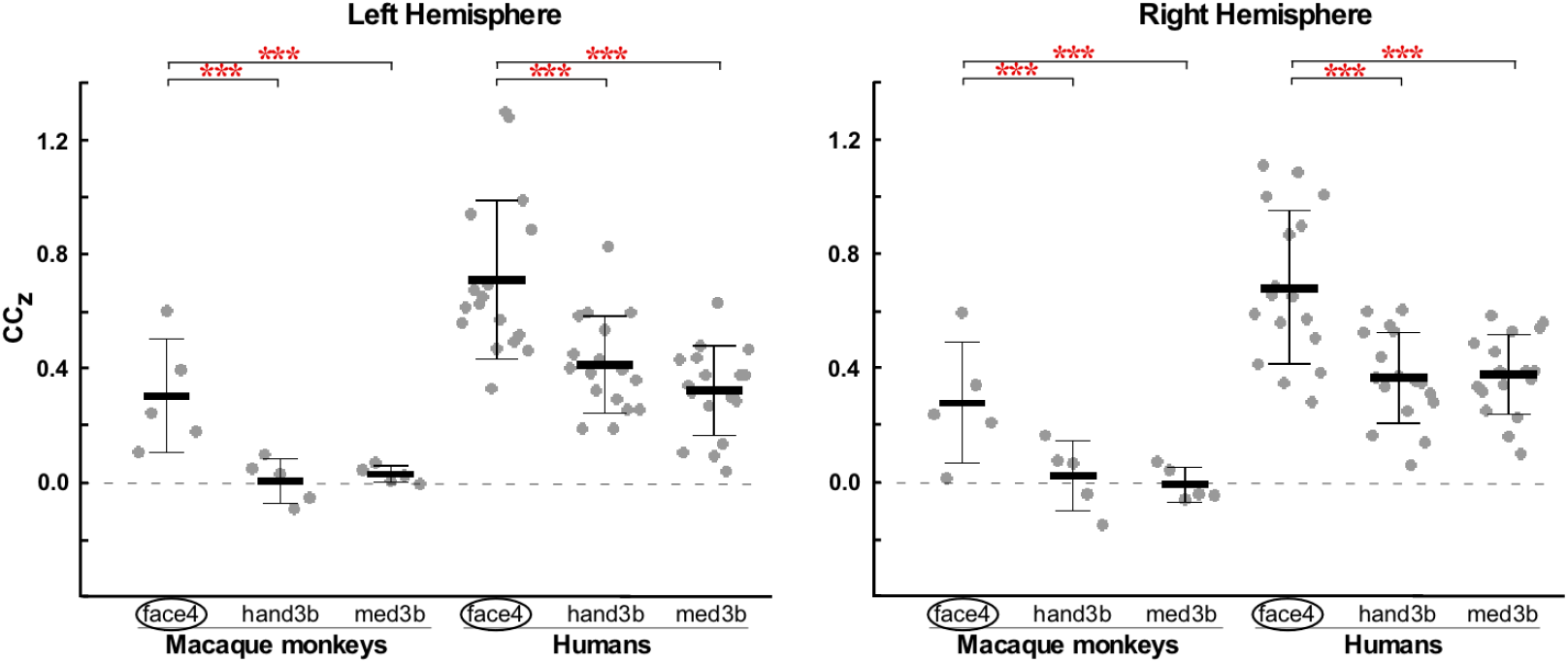
Functional connectivity of somatosensory face representation, face3b with rostrally adjacent homotopic motor representation in area 4 (face4; enclosed in oval) and its comparison with connectivity with other representations (hand3b and med3b) in the same hemisphere of macaque monkeys and humans. Data is shown for both the hemispheres. Each dot on the plot denotes Fisher-z transformed correlation value (CCz) for a single subject in monkeys and humans. Horizontal lines show the mean (thick line) and ± S.D (thin line) of the connectivity strength. Asterisks*** denote statistically significant difference, *p* < 0.0001, two-way ANOVA (main effect of ROI’s); *p* < 0.01, post hoc Tukey test.

In the left hemisphere of both the species, the connectivity of hand3b with hand4 was significantly higher than with face3b or med3b (For LH, macaque, *F*_(2, 81)_ = 22.86, *p* < 0.0001; humans, *F*_(2, 51)_ = 12.05, *p* < 0.0001; two-way ANOVA; *p* < 0.01, *post hoc* Tukey test). In both species, the connectivity of hand3b with hand4 in the right hemisphere was higher, but not statistically significantly when compared to connectivity with face3b or with med3b (For RH, macaque, *F*_(2, 81)_ = 2.37, *p* = 0.10; humans, *F*_(2, 51)_ = 5.93, *p* < 0.004; two-way ANOVA; n.s, *post hoc* Tukey test).

Connectivity of med3b with med4 in humans was significantly higher than with face3b or hand3b, but only in the left hemisphere (*F*_(2, 51)_ = 12.56, *p* < 0.0001; two-way ANOVA; *p* < 0.01, *post hoc* Tukey test). The mean connectivity of med3b with med4 in the right hemisphere of both species was higher than with face3b or with hand3b but the difference was not statistically significant (for RH, macaque, *F*_(2, 81)_ = 0.845, *p* = 0.4334; humans, *F*_(2, 51)_ = 7.55, *p* = 0.0013; two-way ANOVA; n.s, *post hoc* Tukey test).

### 3.3. Comparison of the functional connectivity in monkeys and humans

The functional networks of entire area 3b and different body part representations had many similarities in both macaque monkeys and humans (Fig. 4 and 10). The network if determined for entire area 3b largely represents nodes of individual networks of face3b, hand3b and med3b. In both monkeys and humans, face3b had functional connectivity with regions in the primary motor cortex as well as other areas. Only the face3b was functionally connected to S2 and PMv, bilaterally for humans and in the right hemisphere for monkeys (Fig. 8). The hand3b and med3b of both the hemispheres in both species had network connectivity restricted to area 3b and area 4 (Fig. 6).

**Fig. 10.**
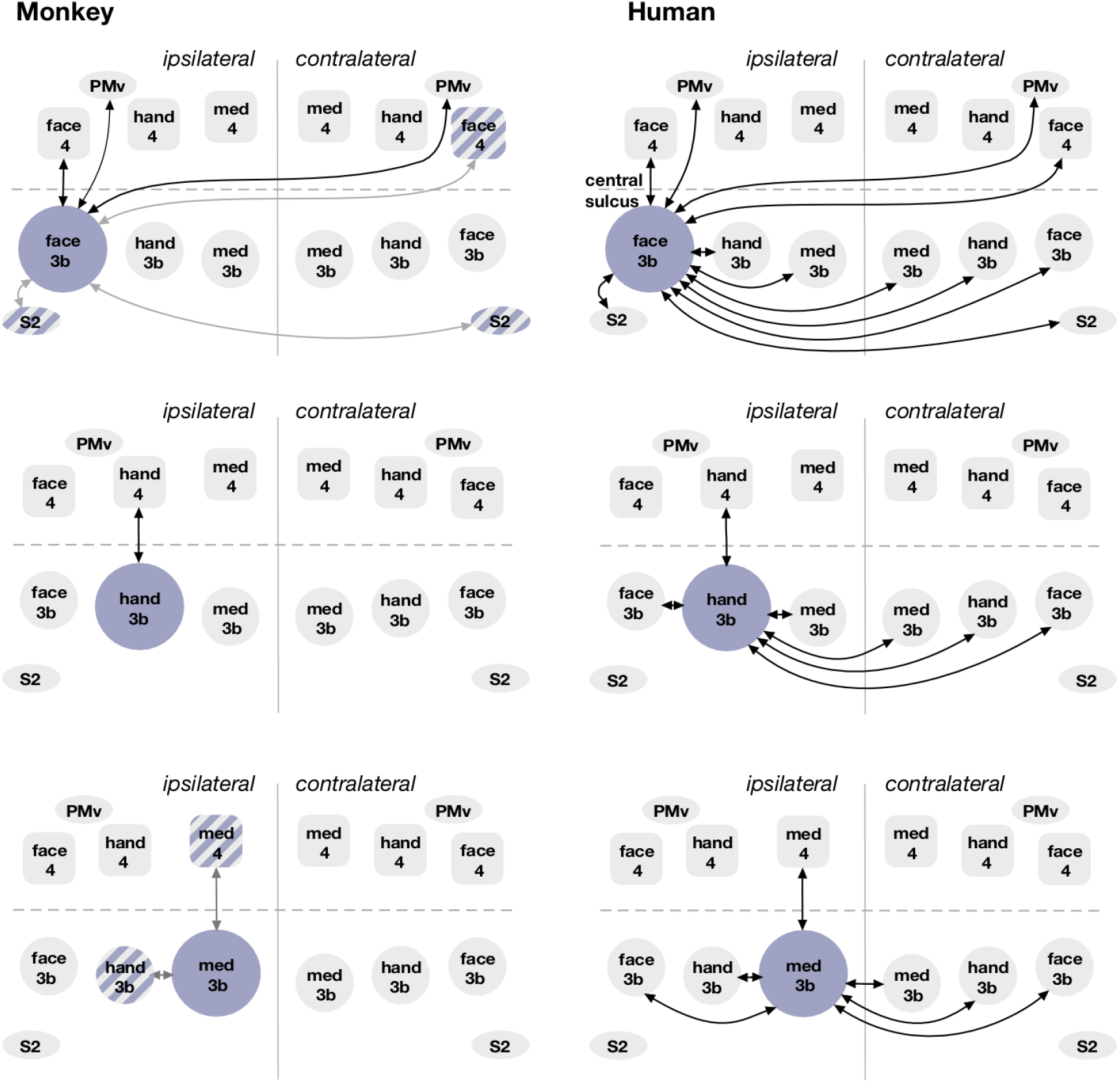
A schematic showing resting-state functional connectivity of different ROI’s - face3b, hand3b, and med3b (large circles) in the ipsilateral and contralateral hemispheres in macaque monkeys (left panels) and humans (right panels). Significant connections common to both hemispheres are depicted. Spheres represent topographic representations in area 3b, rounded squares in area 4; and ovals represent areas other than the primary cortices. Blue shaded regions show areas with statistically significant ROI-ROI correlations as shown by double headed arrows. The regions filled with grey did not have any significant correlations. In addition, significant connections in one hemisphere of monkeys but both hemispheres of humans are shown for comparison by grey lines and hashed fills. One sample t-test, *p* < 0.05 (monkeys), *p* < 0.00001 (humans). Arrows in the schematic do not imply any directionality.

However, there were few notable differences in the functionally connected regions and their connectivity strengths between the two species. Face3b connections in macaque monkeys showed significant bilateral correlations with parietal area 7b and putamen, which were not observed in humans (Fig. 5). The face3b network in monkeys did not show any connections with the regions on the medial wall at the border of area 4 and the supplementary motor area, which was seen in humans (Fig. 5).

Significant connectivity of the body part representations in area 3b to homotopic representations in contralateral area 3b was observed only in humans (Fig. 6 and 10). Finally, the significant connectivity between different body part representations within area 3b i.e. face, hand and medial regions was also observed only in humans (Fig. 6 and 10).

## 4. DISCUSSION

We performed seed-based correlation analysis to determine resting-state connectivity of different body part representations in area 3b of macaque monkeys and humans. Previously both somatosensory and motor areas have been lumped together for connectivity analysis. Our results show that there is a characteristic somatosensory network for each body part representation, which is largely similar in macaque monkeys and humans. Some of the main observations from our ROI-ROI and ROI-voxel analyses are discussed below. The results are summarized in Figure 10.

### 4.1 Understanding brain connectivity using resting-state correlations

Resting-state functional connectivity networks, which are determined from the time-series correlation of different brain regions in a task free condition have been described for different mammalian species including humans, monkeys, cats, ferrets, rats and mice (Biswal et al., 1995; Lu et al., 2012; Popa et al., 2009; Stafford et al., 2014; Vincent et al., 2007; Zhou et al., 2016). Resting-state networks have been used to delineate functional brain networks such as somatomotor, visual and auditory networks, the dorsal and ventral attention systems, the fronto-opercular salience region, and what is known as default mode network (Biswal et al., 1995; Cordes et al., 2000; De Luca et al., 2005; Fox et al., 2006; Fox et al., 2005; Greicius et al., 2003; Lowe et al., 1998; Raichle et al., 2001; Seeley et al., 2007). The resting-state networks also predict brain areas that would be active during stimulus driven activation and during various cognitive tasks (De Luca et al., 2005; Hampson et al., 2006; Seeley et al., 2007; Tavor et al., 2016; Vincent et al., 2006).

We determined resting-state network connectivity for different topographic representations in the somatosensory cortex of macaque monkeys and humans. Previous reports in macaque monkeys described the resting-state network considering all the somatomotor areas as a single ROI (Hutchison et al., 2011), or at the most after dividing the somatomotor areas into dorsal and ventral subdivisions (Mantini et al., 2013). We show here that each body part representation in area 3b has its own distinct network, which is different from that for other body parts. Network of the entire area 3b largely reflects sum of all the individual networks. The somatosensory network for complete area 3b described here is generally similar to as those for the ‘somatomotor networks’ described in previous studies (Biswal et al., 1995; Kuehn et al., 2017; Mantini et al., 2013; Vincent et al., 2007). However, some of the nodes such as area 5, 7, putamen, and insula are revealed only when area 3b was considered separately as seed ROI, suggesting importance of area specific ROI’s for connectivity analysis.

Previously, neuroimaging studies have divided area 3b into two independent networks - a ventral network that comprises of the tongue and possibly lower face representation, and a medial network comprised of the hand and other body part representations (Power et al., 2011; Yeo et al., 2011). These were defined by network parcellation. We show here that the face representation, even after excluding the intraoral structures is an independent network separate from the ‘hand network’. It is likely that the face and oral structures would form separate networks, given they have different connectivity (Cerkevich et al., 2014; Iyengar et al., 2007) and are revealed as separate myelin rich modules in histological preparations of the monkey cortex (Jain et al., 2001).

We predict that if we determine functional connectivity of cortical representations at higher granularity, more details of connectivity will be revealed. For example, each digit might reveal a specific connectivity due to digit specific, and divergent and convergent connections (Ashaber et al., 2014; Negyessy et al., 2013; Wang et al., 2013). At an even higher resolution different parts of the digits, e.g. proximal and distal likely reveal different connectivity (Liao et al., 2013). Furthermore, each of the body part representation in our med3b will likely have different connectivity pattern (Krubitzer and Kaas, 1990).

### 4.2 Functional connectivity reflects anatomical connections

The resting-state functional networks reflect anatomical connections as determined using neuroanatomical tracers or lesion studies. Relationship between functional connectivity and direct anatomical connectivity has been emphasized (van den Heuvel et al., 2009; Wang et al., 2013). While anatomical connections in monkeys have been described in detail, information on the anatomical connectivity in humans is sparse and generally indirectly inferred. Our results show that in monkeys area 3b has connectivity with areas 1, 2, 5, 7, S2/PV, PMv and insula, the areas that are known to have anatomical connections with area 3b (Burton et al., 1995; Darian-Smith et al., 1993; Jones and Powell, 1969a; Pons and Kaas, 1986). Differences in the ipsilateral and contralateral resting-state connectivity observed for different body part representations - face3b, hand3b, and med3b reflects differences in the underlying anatomical connectivity of these representations (Jones and Powell, 1969b; Killackey et al., 1983). For example, we found that face3b network has the largest number of bilateral nodes which include face4, S2/PV, PMv and insular cortex, and the contralateral face3b. This reflects anatomical connectivity of face representation in area 3b. Face representation in area 3b is known to receive direct projections from area 3a, 1, 2, S2/PV, PMv and Insula (Cerkevich et al., 2014; Disbrow et al., 2003).

Connectivity of area3b is also seen with areas 3a, 1 and 2 (see Fig. 4 and 5) but is not separately analyzed here because of difficulties in placing ROI’s that are clearly distinct from area 3b.

In macaque monkeys, the face representation also showed significant correlation with the posterior parietal area 7b and putamen in the ROI to voxel analysis. Neuroanatomical studies have shown connectivity of the face representation in area 3b with the rostro-lateral part of area 7b (Burton et al., 1995; Lewis and Van Essen, 2000), and large parts of area 7b has somatosensory responses to mouth and face (Hyvarinen, 1981). In monkeys topographic projections from the somatosensory cortex to putamen have been seen using anatomical tracers (Jones et al., 1977; Kunzle, 1977) and in lesion studies (Kemp and Powell, 1970). The voxels in the lateral putamen correlating with face3b are in a similar location as the somatosensory projections to putamen (Kunzle, 1977).

As compared to face3b, the network of other ROIs i.e., hand3b and med3b had fewer nodes, which were largely confined to the primary motor cortex (see Results, and Fig. 10). In monkeys, hand3b did not functionally connect to the hand representation in contralateral area 3b. This reflects the lack of homotopic callosal connections of the hand representation in area 3b (Killackey et al., 1983). In monkeys, bilateral correlation of hand3b with area 5 was observed. In area 5 there are converging inputs in the forelimb representation with neurons having large receptive fields on the arm and the hand (Padberg et al., 2005; Seelke et al., 2012).

Differences in the functional connectivity of the face and hand representations in area 3b with PMv also reflect differences in the anatomical connectivity. Although the strongest projections from area 3b to PMv are from the face representation, PMv also has minor inputs from the hand representation (Dancause et al., 2006). In humans we observed hand3b to PMv connectivity in the left hemisphere.

In both species, the bilateral connectivity of area 3b representations to homotopic representations in the contralateral hemisphere is in agreement with the known callosal connectivity of body part representation in area 3b, where face and trunk representations have more callosal connections as compared to the hand representation (Killackey et al., 1983; Krubitzer and Kaas, 1990; Pandya and Vignolo, 1969). Moreover, the midline body parts such as the face and the trunk have occasional bilateral receptive fields in area 3b (Conti et al., 1986; Dreyer et al., 1975; Eickhoff et al., 2008; Fabri et al., 2005). Although most of these seems to be contributed by callosal connections (Fabri et al., 2006) peripheral contribution cannot be ruled out (Iwamura et al., 2001). These peripheral bilateral inputs can also possibly contribute to interhemispheric connectivity. Thus, the functional connectivity of different body part representations reflects anatomical connectivity.

However, we did not observe any significant resting-state correlations of hand representation in area 3b with S2 in spite of their direct anatomical connectivity (Eickhoff et al., 2007; Krubitzer et al., 1995). Among the five monkeys scanned, the hand3b-S2 correlation was found only in two animals using seed-to-voxel analysis at lenient thresholds and does not emerge as significant correlation in the ROI-ROI analysis. In humans also, the hand3b had correlations to S2 at lenient thresholds observed only in the seed-to-voxel analysis.

### 4.3 Functional connectivity is not always dependent upon direct structural connectivity

It has been suggested that functional connectivity is not necessarily constrained by absence of direct structural connectivity (Raichle, 2015). The functional connectivity networks are not only driven by monosynaptic connections but also polysynaptic connections (Honey et al., 2009; Vincent et al., 2007). Thus resting-state coherence observed between areas that lack monosynaptic connections likely reflects emergent cortical network properties (Adachi et al., 2012; Vincent et al., 2007). Our results also reveal this aspect of structure-function relationship of functional networks. For example, we found strong functional association between area 3b and area 4 representations, which lack direct anatomical connections. These functional correlations might reflect the indirect anatomical links through somatosensory area 2 and 5 which receive projections from area 3b and send projections to the motor cortex (Darian-Smith et al., 1993; Jones et al., 1978; Vogt and Pandya, 1978).

The functional connectivity between representations across the central sulcus points to information flow between homologous representations in area 3b and area 4 required for coordinated activity in these brain areas required for sensorimotor tasks. There is also a phylogenetic closeness of the neuronal mass populating the similar body representations that mediate coordinated neural activity in the sensory and motor regions (Flechsig, 1920 as cited in Kuehn et al., 2017). A functional association between motor and somatosensory areas has been observed as spatiotemporal coherence in local field potential (LFP) signals (Arce-McShane et al., 2016; Murthy and Fetz, 1992).

There are other examples illustrating that resting-state functional connectivity also reflects indirect or higher order anatomical connectivity. For example, connectivity of area 3b with insular cortex is likely a reflection of indirect anatomical connectivity via the secondary somatosensory cortex (Burton et al., 1995; Friedman et al., 1986). Functionally, innocuous somatosensory stimulation of face is known to elicit neuronal responses in the granular regions of insula (Schneider et al., 1993).

### 4.4 Resting networks of area 3b are similar in humans and monkeys

Although separated by 25 million years of evolution (Kaas, 2004, 2012), macaque monkeys and humans share similarities in the structural connectivity and functional networks of different brain areas (Goulas et al., 2014; Mantini et al., 2013; Sallet et al., 2013). The somatomotor networks have high topological correspondence in humans and monkeys (Mantini et al., 2013). The current study is the first to determine and directly compare resting-state networks of different topographic representations in both humans and macaque monkeys. We used somatotopy specific seed ROIs that were also validated using localizer scans in both macaques and humans, assuring accuracy of the observed networks.

Our results show that connectivity of different body part representations in area 3b viz. face3b, hand3b and med3b is similar in monkeys and humans (see Fig. 5, 6). These similarities include (1) face 3b having widespread functional connectivity with networks that include ipsilateral face4, bilateral PMv, and the insular cortex, and (2) functional connectivity of the face and hand representations in area 3b with homotopic representations in area 4. These similarities likely reflect many behavioral similarities between the monkeys and humans. Both these species are highly social with a variety of complex facial expressions that are important for maintaining the social structure (Burrows, 2008). The widespread functional associations seen especially for the face representation in both monkeys and humans might underlie the important role of facial structures in verbal and nonverbal communication which enables the rich socio-cultural lives of primate communities. Although the difference in the ability to use the hand vary considerably, monkeys are able to make a large variety for grasps just like humans (Macfarlane and Graziano, 2009). Both these species have opposable thumbs enabling fine grasping ability (Marzke, 1997).

Previous neuroanatomical data from monkeys has shown that connectivity between different areas is strongest between homotopic body part representations (Ashaber et al., 2014; Negyessy et al., 2013; Wang et al., 2013). Our results also show stronger functional connectivity between homotopic body part representations, thus reinforcing the importance of somatotopic representations in information processing.

There were also certain differences in the functional connectivity networks of two primate species (see Fig. 5, 10), probably reflecting species specific behavioral differences. Many functional correlations were found only in humans. For example, humans showed correlation of face3b with voxels at M1/SMA border on the medial wall while the monkey brains did not. Bilateral connections between homologous area 3b representations were observed only in humans; as well as the significant connectivity between face, hand and the medial regions of area 3b was present only in humans. The comparatively higher connectivity of face representation in humans could be a reflection of the importance of facial gestures for emotions and social cues. The bilateral connectivity of the somatosensory representations in humans reflects a wider repertoire of behavior in humans that uses bilateral coordination in bipedal humans including their complex object manipulation abilities that requires fine tactile inputs.

### 4.5 Limitations of the study and methodological considerations

The difference in functional connectivity between humans and monkeys could reflect evolutionary differences between the two species. However, there are technical considerations that should also be taken into account while interpreting the results.

The inter-species differences in functional connectivity could be due to differences in the brain organization i.e. the number and sizes of the cortical areas, their specialization, and anatomical connectivity. For example, correlations of area 3b representations with putamen, area 7 and area 5 were found only in macaque monkeys. These, differences in functional connectivity might also be due to the differences in brain sizes of the two species which can give rise to alterations in neuronal wiring and information processing networks (Kaas, 2000).

The mean correlation strengths of connections in monkeys were always lower than in humans (e.g. Figs 7, 8 & 9). Larger size of human brain along with its dense white matter fiber connections can give rise to stronger correlation values.

However, the correlation strength difference could also be due to differences in the sizes of the monkey and human brain scanned in the same magnet. Although we used a knee coil for monkeys to improve the filling factor, the signal to noise ratio was better for the human brain.

Difference in the brain states can also result in the difference between the human and monkey data. We scanned monkeys in anesthetized state for ease of handling, while human participants were awake. Because of the anesthetized state of monkeys, there could be a reduction in the BOLD signal strength and consequently reduction in the calculated correlation strengths (Bettinardi et al., 2015; Grandjean et al., 2014; Vincent et al., 2007). Isoflurane, used in the current study, is known to depress the brain activity and cause general nonselective suppressive effects on local functional connectivity of fine-scale cortical circuits in a dose dependent manner (Hutchison et al., 2014; Wu et al., 2016). Thus, the functional connectivity determined during the awake state might be reflected as widespread interareal connectivity only in humans, but could become more restricted in anaesthetized monkeys.

Different anesthetic agents affect the connectivity patterns differently (Paasonen et al., 2018). Ketamine, an NMDA antagonist, has been shown to reduce the intrinsic brain connectivity in primary somatosensory and auditory cortices (Niesters et al., 2012). We chose isoflurane because comparative studies using different anesthetic agents show that isoflurane is an ideal candidate for resting-state connectivity studies in animal models (Grandjean et al., 2014; Jonckers et al., 2014; Lv et al., 2016). Moreover, the isoflurane levels in the current study (0.8% maximum) were lower than the most resting-state studies which use up to 1.25%-1.5% isoflurane (Hutchison et al., 2014; Vincent et al., 2007; Wu et al., 2016). Although the amplitude of functional correlations might reduce under anesthesia, the conservation of functional networks across different brain states such as arousal and sleep have been described (Fukunaga et al., 2006; Hutchison et al., 2013). This suggests that despite the differences in the brain state while scanning, the differences observed between the two species could be actual species-specific differences.

Although there are reports that the functional connectivity is not static but dynamically variable across different time scales (Chang and Glover, 2010; Hutchison et al., 2013), we used averaged connectivity measures from each ROI for the analysis. Certainly, different analysis methods considering non-stationary transient state changes in connectivity can give more complete information regarding the baseline spontaneous activity in the brain. However, the topographic variation of resting functional networks across different body part representations was visible even while using zero-lag, time averaged correlation measures. Despite these limitations we found remarkable similarities between humans and monkeys.

## 5.0 CONCLUSIONS

Our results show that the network architecture described as ‘somatomotor network’ is a composite structure and is comprised of multiple independent and different networks of different topographic representations. Care needs to be taken while assigning the network characteristics uniformly to the participating nodes. A similar recent study has described the non-uniformity of the task negative default mode network (DMN) in humans and has found that subnetworks exist within the DMN (Braga and Buckner, 2017). Previous reports on the intrinsic functional connectivity of human brain discuss the intra network heterogeneity within somatomotor network and suggest the importance of topographic areas in delineating network boundaries (Kuehn et al., 2017; Long et al., 2014; Power et al., 2011; Yeo et al., 2011). The current study extends the understanding of the normal structure of spontaneous connectivity and describes how it varies across different body part representations in somatosensory area 3b in macaque monkeys and humans. In all the acquired fMRI sessions in both the species, the somatosensory ROIs consistently described distinct functional subnetworks, which largely reflected the underlying anatomical connectivity patterns. The results suggest that rather than considering the entire ‘somatomotor’ area as a whole, connectivity network analysis should take body part representations into consideration. This knowledge of the somatotopy dependent connectivity is crucial in understanding changes in the information processing within these networks in disease and injury conditions such as spinal cord injury that cause large-scale somatotopic reorganization.

## ACKNOWLEDGEMENTS

Authors are grateful for funding by Department of Biotechnology, Government of India to N.J. (No.BT/PR7180/MED/30/907/2012) and core funds from National Brain Research Centre.We also thank Mr. Atanu Datta, Mr. Arun E V R and Mr. Raghav Shankar for assistance with acquisition of monkey fMRI data, and Mr. Jitender Ahlawat for technical assistance. We thank Dr. V Rema for helpful comments on the manuscript.

